# Schrödinger’s phenotypes: herbarium specimens show two-dimensional images are both good and (not so) bad sources of morphological data

**DOI:** 10.1101/2020.03.31.018812

**Authors:** Leonardo M. Borges, Victor Candido Reis, Rafael Izbicki

## Abstract

1. Museum specimens are the main source of information on organisms’ morphological features. Although access to this information was commonly limited to researchers able to visit collections, it is now becoming freely available thanks to the digitization of museum specimens. With these images, we will be able to collectively build large-scale morphological datasets, but these will only be useful if the limits to this approach are well-known. To establish these limits, we used two-dimensional images of plant specimens to test the precision and accuracy of image-based data and analyses.
2. To test measurement precision and accuracy, we compared leaf measurements taken from specimens and images of the same specimens. Then we used legacy morphometric datasets to establish differences in the quality of datasets and multivariate analyses between specimens and images. To do so, we compared the multivariate space based on original legacy data to spaces built with datasets simulating image-based data.
3. We found that trait measurements made from images are as precise as those obtained directly from specimens, but as traits diminish in size, the accuracy drops as well. This decrease in accuracy, however, has a very low impact on dataset and analysis quality. The main problem with image-based datasets comes from missing observations due to image resolution or organ overlapping. Missing data lowers the accuracy of datasets and multivariate analyses. Although the effect is not strong, this decrease in accuracy suggests caution is needed when designing morphological research that will rely on digitized specimens.
4. As highlighted by images of plant specimens, 2D images are reliable measurement sources, even though resolution issues lower accuracy for small traits. At the same time, the impossibility of observing particular traits affects the quality of image-based datasets and, thus, of derived analyses. Despite these issues, gathering phenotypic data from two-dimensional images is valid and may support large-scale studies on the morphology and evolution of a wide diversity of organisms.

## 1 Introduction

Specimens held in natural history collections are our primary source of information on the diversity, morphological variety, and spatial and temporal distribution of living creatures. Based on this information, we obtain knowledge on many areas, such as environmental changes, public health, and evolution (Suarez and Tsutsui, 2004; Law and Salick, 2005; Babin-Fenske et al., 2008; Davis et al., 2015). Until recently, however, only people with access to museum collections could study these specimens. Now, almost all institutions are making their specimens more accessible through digitization.

Digitization methods vary as much as the biodiversity included in museums. Zoological collections may require the use of complex methods that better capture animal form, such as computer tomography, magnetic resonance imaging, or photogrammetry (e.g. Berquist et al., 2012; Falkingham, 2012; Keklikoglou et al., 2019). 2D imaging (photographs or flat scans), which has also been used for animals (e.g. Mantle et al., 2012; Schmidt et al., 2012), is particularly useful for flat specimens. Indeed, the flat, regular in size, and easy to image nature of plant samples has led to millions of plant specimens being available online (Le Bras et al., 2017; Soltis, 2017), and ready to be used as part of scientific research.

Taxonomy is one area in which digitization has been very fruitful. For example, automatic species identification software can substantially speed up taxonomy and specimen curation (Wang, Ji, Liang and Yuan, 2012; Remagnino et al., 2016; Carranza-Rojas et al., 2017). These tools, however, usually classify specimens by their overall image patterns, without making assumptions about organ identity (Wang, Ji, Liang and Yuan, 2012; Favret and Sieracki, 2016; Remagnino et al., 2016). Trait delimitation with images and derived data extraction (Corney, Tang, Clark, Hu and Jin, 2012; Corney, Clark, Tang and Wilkin, 2012; Wang, Lin, Ji and Liang, 2012; Gehan et al., 2017; Martineau et al., 2017), although promising, is still complex and underdeveloped. Even if we still have to wait for automatic data extraction to become highly efficient, imaging specimens facilitates data acquisition by researchers themselves, particularly if used in collective efforts to build comprehensive morphological datasets (e.g. O’Leary et al., 2013).

Such phenotypically diverse datasets may support advances in comparative and evolutionary biology (Laing et al., 2018). However, in comparison to genomic data, they take more time and money to produce (Burleigh et al., 2013). Tools that use specimens images are one option to overcome these challenges (Burleigh et al., 2013). However, this approach has limits. First, image resolution may limit observations, especially of small traits. Second, the nature of organisms themselves, or of specimen preparation, hides some features. For example, stamens in beans flowers are naturally hidden by the petals, and the wings of pinned butterfly specimens usually cover their legs. Although this is not a problem when one has access to specimens, it may become an issue for images. Thus, such variations in the preparation and nature of specimens may impact image-based data collection.

To evaluate the influence of these restrictions on the acquisition of morphological data from digitized specimens, we asked (1) if image measurements significantly differ from specimen measurements, and (2) if image-based datasets differ from the ones built with specimens to the point of affecting morphological analyses. To answer our first question, we compared measurements taken from herbarium specimens and their two-dimensional digital image. To answer our second question, we compared the results of multivariate analyses of plant specimens with analyses simulating image usage.

## 2 Materials and Methods

### 2.1 A word on terminology

Before outlining our analyses, we explain some of the terms we use, in particular accuracy and precision (for details, see Streiner and Norman, 2006). We consider precision to be an estimate of the variation between multiple measurements of the same feature. If values for these measurements do not differ widely, precision is high. Accuracy is treated as the difference between a measurement and a reference value and also applied here to judge the results of multivariate analyses. A close similarity between the measurement (or results) and the reference value indicates high accuracy. Because our goal is to evaluate the confidence of images as a data source in comparison to that obtained from specimens, we treated specimen data as the reference values against which we judged the accuracy of image data.

### 2.2 Measurement precision and accuracy

To test if measurements made from images and specimens differ, we measured different leaf parts of specimens belonging to five species (hereafter, `measurement dataset`). These species (*Inga vera* Willd., *Ocotea divaricata* Mez, *Piper anisum* (Spreng.) Angely, *Smilax fluminensis* Steud. and *Sphagneticola trilobata* (L.) Pruski) were selected to encompass different leaf morphologies, such as simple, compound, and lobed leaves with different base and apex shapes. Besides, by using these taxa, we avoided sampling size issues, as they are well represented in the Rio de Janeiro Botanical Garden herbarium (RB; acronym according to Thiers, 2020, continuously updated. See the supplementary material for information on measurements, sampling, and vouchers. Metric data is also available from MorphoBank project 3764 http://morphobank.org/permalink/?P3764). We used a digital caliper to measure specimens and the FSI Viewer v. 5.6.6 software (available from RB’s website; http://jabot.jbrj.gov.br) to measure images of the same leaves from the same specimens. All measurements, both from images and specimens, were made twice to allow the following analyses.

First, we tested tested the precison of measurements made either from the specimens or from the images. To do so, we evaluated how the two repeated measurements are close to each other with the intraclass correlation coefficient (ICC) (Bartko, 1966). Then, we tested the hypothesis that images and specimens have the same average precision. For that, we compared the difference between the two repeated measurements made from images with those made from specimens with a paired t-test. The last test in this context evaluated if precision varies according to measurement scale. For that, we first divided the absolute difference between the two repeated measurements by their average to obtain measurements relative precision. Then, we plotted the relative precision of each observation against a scale reflecting overall trait size.

Having tested for precision, we computed the average of the two measurements of each variable. We used these averages to avoid bias in the following accuracy analyses. For each specimen and variable, we evaluated accuracy between measurements in the images to the specimens by testing their similarity with ICC (Bartko, 1966). We further evaluated accuracy testing the hypothesis that the median difference between image and specimen measurements is zero with a Wilcoxon signed-rank two-sample test (Wilcoxon, 1945). Finally, to quantify the relationship between measurement error and the magnitude of each variable, we plotted the relative error of the measurements (i.e., the absolute difference between image and specimen values divided by image mensuration) against the image measurement itself.

### 2.3 Accuracy of image-based datasets

To test if image-based data impacts analyses of morphological data, we compiled datasets of morphometric studies of different plant groups (Alcantara et al., 2013; Ames et al., 2008; Andres-Sanchez et al., 2009; Bello et al., 2018; Bünger et al., 2015; Egan, 2015; Kučera et al., 2006; Marhold, 1992; Poulsen and Nordal, 2005; Rose and Freudenstein, 2014; Slovák et al., 2012; Trovó et al., 2008). These datasets (hereafter `original datasets`) were used to generate three `synthetic datasets`, which were modified to include variation as if they had been compiled from images: the first included bias observed in the measurement accuracy analysis described above; the second included missing data; and the third included both measurement bias and missing data. We generated `synthetic datasets` with the following procedures.

First, we added noise to the `original dataset` using a zero-mean Gaussian distribution with varying values of variance. To do so, we modeled how the standard deviation of the measurements’ error varies as a function of each variable’s magnitude (See supplementary material). We then used this model to simulate different values of standard deviations for the Gaussian distribution of each variable, as to reproduce the same level of measurement error expected for that variable.

Second, to simulate missing observations, we first checked which variables could be obtained from images on a set of up to ten digitized specimens from the same taxa of each morphometric study (see supplementary material for information on variables and vouchers). Using this information, we modeled patterns of missing data, which we used to mask observations in the `original dataset`, producing synthetic data including missing observations (Fig. 1). Finally, we combined the approaches just described to simulate the presence of both noise and missing data.

**Figure 1:**
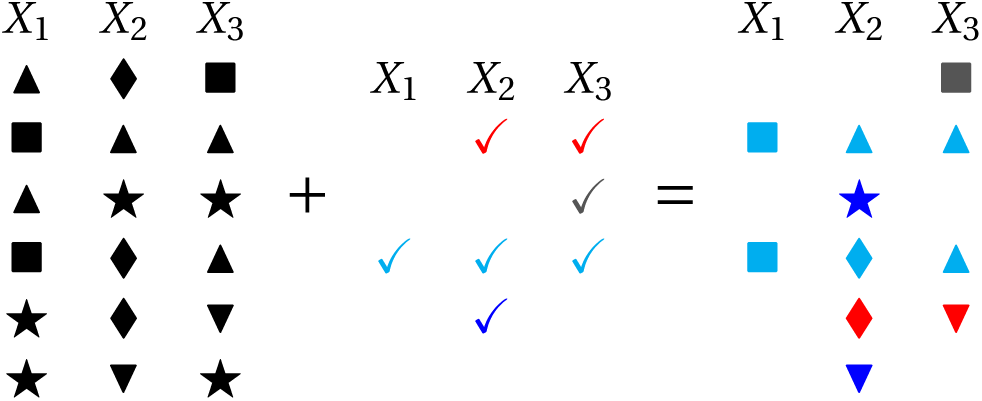
Illustration of the masking procedure. Left: `original` data set. Center: missing patterns obtained using digitized specimens; question marks indicate missing information and check-marks indicate available variables. Right: masked data used for constructing the `synthetic dataset`, obtained by applying a randomly chosen missing pattern (from the middle table) to each row of the original data set.

With `original and synthetic datasets` at hand, we compared agreement between results of Principal Component Analyses (PCA) of all of them. Besides being one of the most common methods in morphometric analyses, differences in PCA results summarize the overall variation of datasets being compared. Thus, PCA comparison captures differences between data acquisition methods (images vs. specimens).

Our protocol first used `pcaMethods` (Stacklies et al., 2007) to run a Probabilistic PCA (Roweis, 1998) for each dataset, which automatically handles missing data with an expectation-maximization algorithm. We then measured the agreement between the first four principal components (PC) inferred for `original` and `synthetic data` by computing Pearson’s correlation coefficient (Pearson, 1920) between the scores obtained for each observation. High correlation indicates agreement between analyses, and consequently between acquisition methods. However, as PC correlations may not capture overall dataset variation (Yang and Shahabi, 2004; Melo et al., 2015), we also used `EvolQG` (Melo et al., 2015) to measure PCA similarity (Yang and Shahabi, 2004) between `original` and `synthetic data`. PCA similarity weights PC correlation by their eigenvalues (Yang and Shahabi, 2004; Melo et al., 2015), and, thus, takes into account how much data variability is expressed in each PC.

The process described above was repeated 100 times (that is, 100 `synthetic datasets` were created), so we could evaluate the accuracy of image-based analyses to specimen-based PCAs with mean values and 95% confidence intervals.

## 3 Results

Measurements taken both from images and specimens have high internal precision, as seen in intraclass correlation coefficient values of 100.0%. Moreover, the hypothesis that precision is the same between images and specimens was not rejected (p=0.678). Indeed, Fig. 2 indicates that measurements made on images and specimens have similar precision, and that, independently of the source, the precision decreases together with trait size.

**Figure 2:**
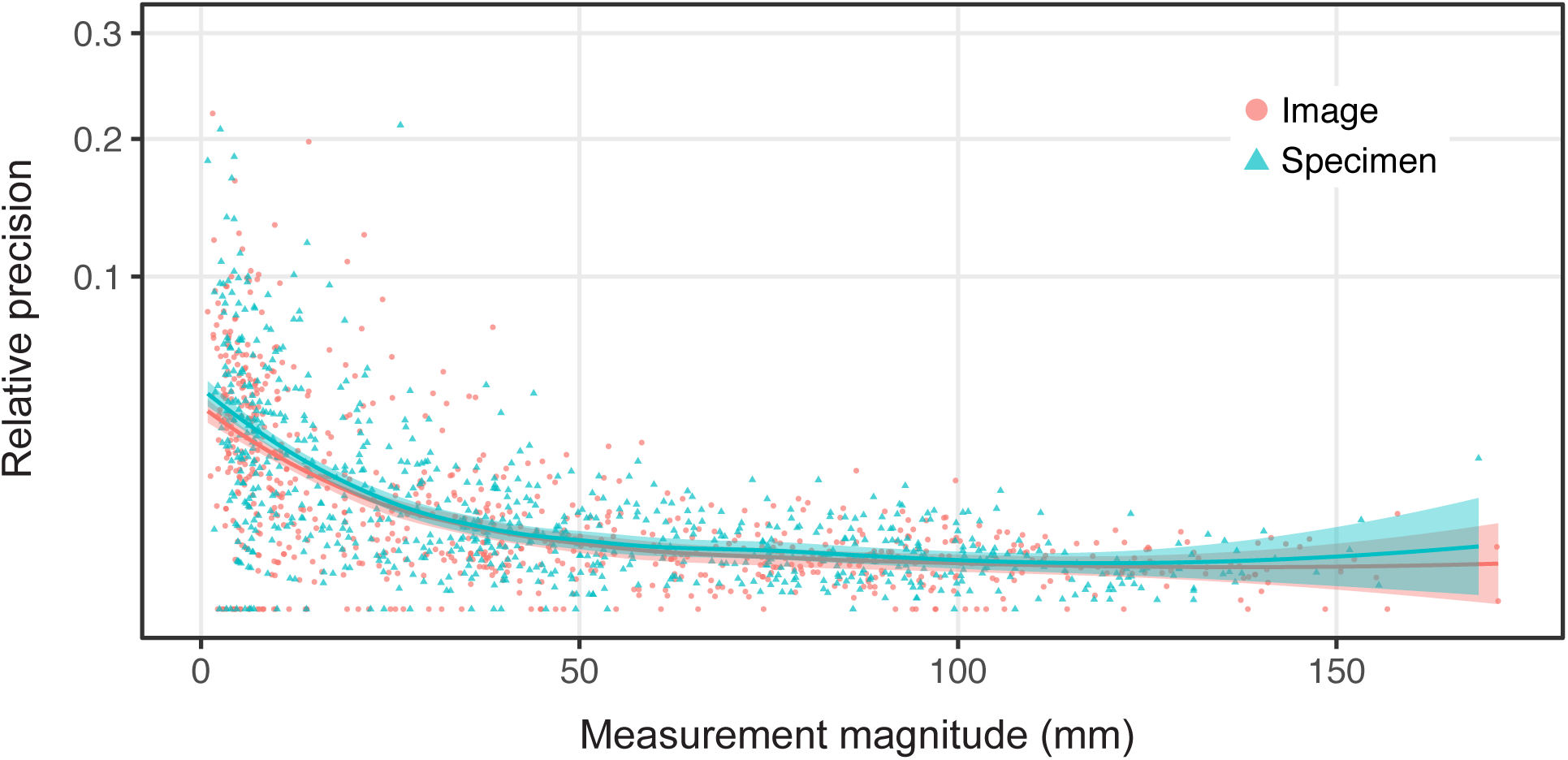
Relationship between the relative precision of measurements (i.e., the absolute difference between two repeated measurements divided by their average) and magnitude of each variable. Dots are different variable observations from the `measurement dataset`, either made on images (red) or on specimens (blue). The curves indicate the mean trend, along with their standard errors. *Y* -axis displayed in square root scale.

Similarly, high ICC values indicate that measurements taken from images are accurate overall (Fig. 3). This lack of bias is reinforced by non-rejection of the hypothesis that they would not differ from specimens measurements (Fig. 4). At the same time, although measurements of larger traits are quite similar between specimens and images, measuring smaller organs with images is not as accurate. For example, blade length measurements of *Ocotea divaricata* or *Piper anisum*, which vary between 50– 160 mm, are extremely similar between specimens and images, as shown by the almost perfect fit of points to the identity line (curve *y = x*) in Fig. 3. On the other hand, points comparing petiole length measurements of the same species (2–15 mm long) are more dispersed around the identity line. This effect is clearly seen in the plot between relative error and specimen measurement values (Fig. 5), which shows that differences between image and specimen measurements are higher for smaller traits (below 30–40 mm).

**Figure 3:**
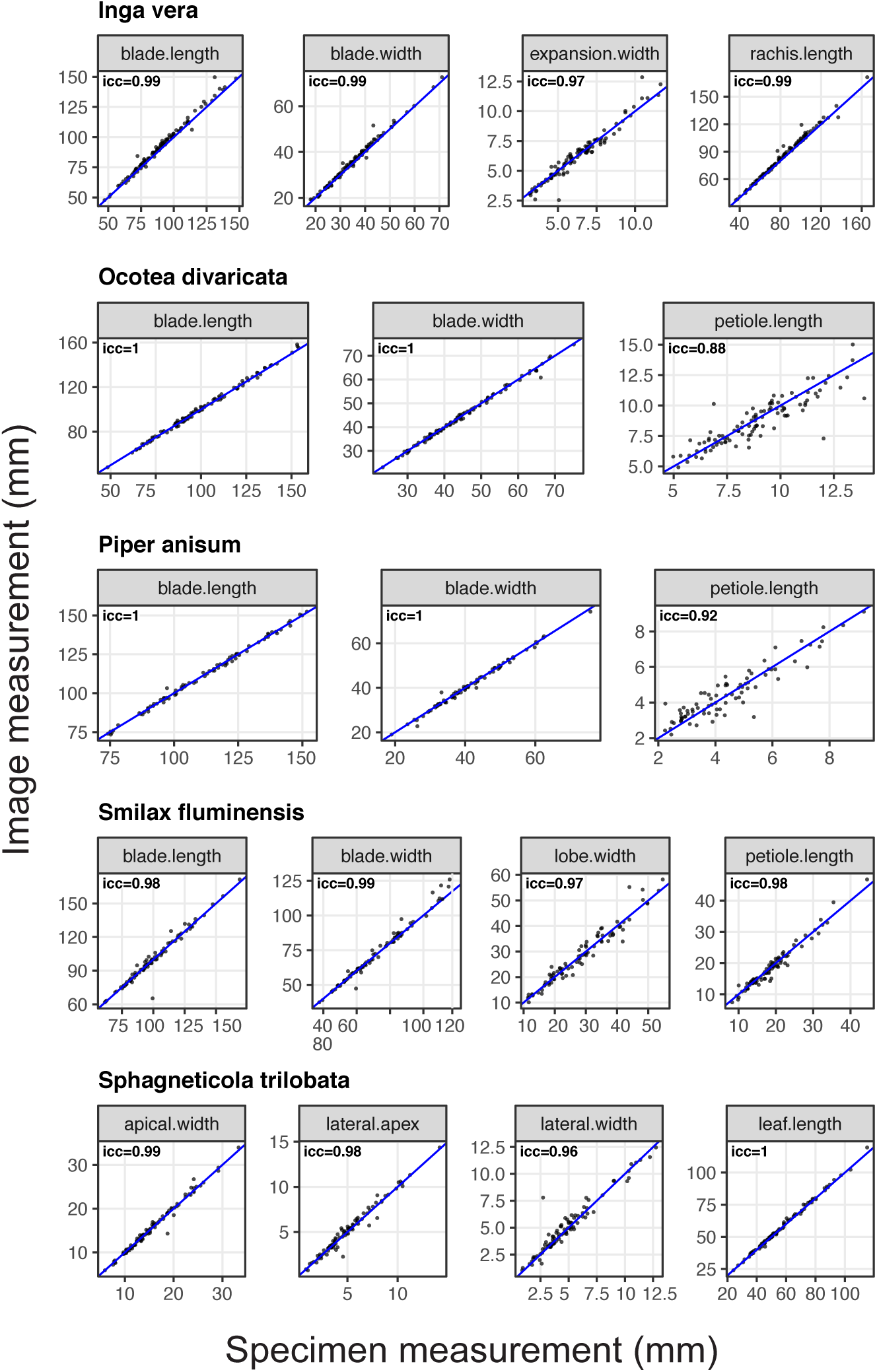
Scatter plots with measurements made from specimen versus measurements made from the images on the `measurement dataset`. Blue lines indicate the line *y = x*, in which both measurements are the same. Each panel also shows the Pearson’s correlation coefficient between the measurements.

**Figure 4:**
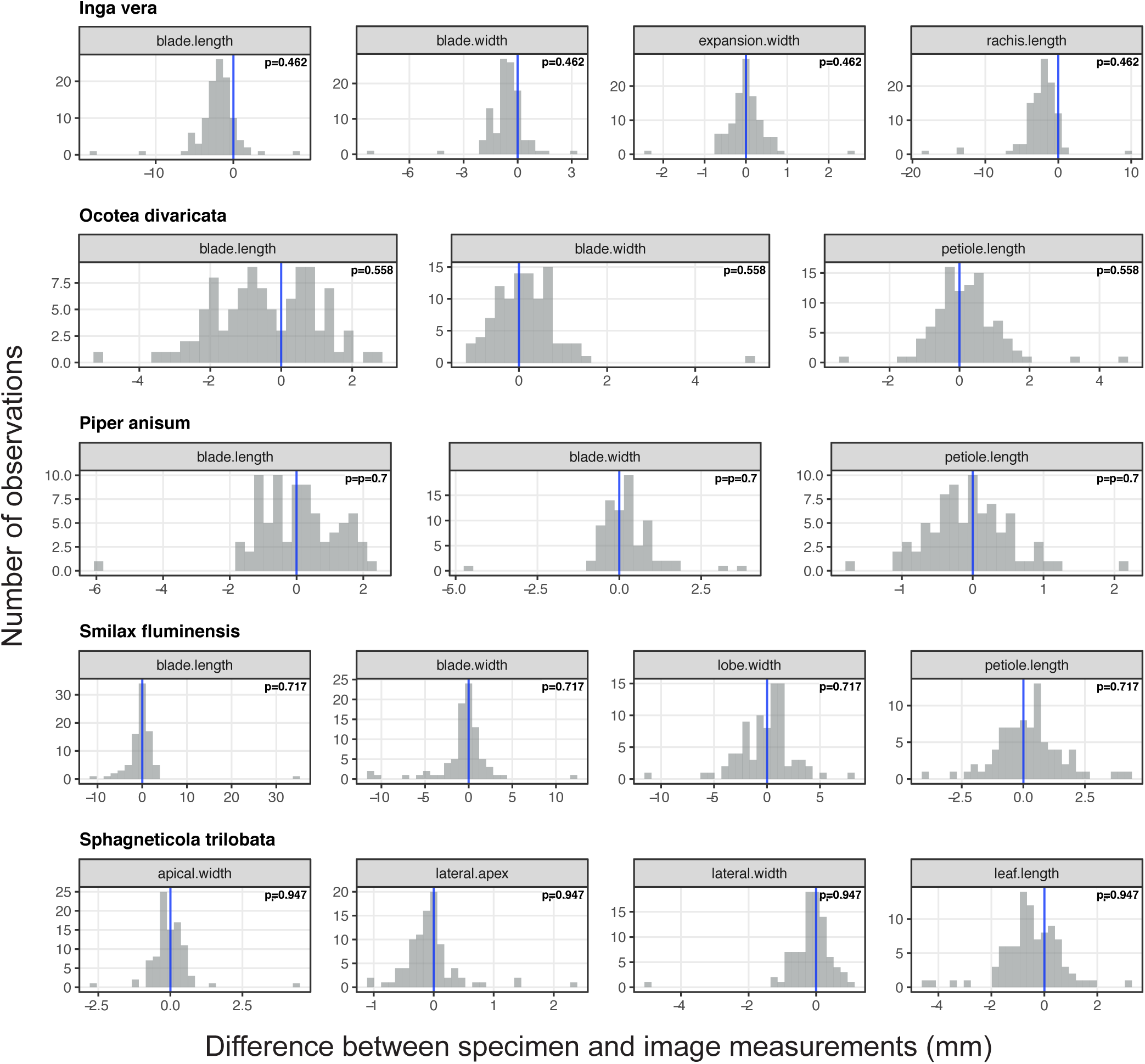
Histogram of the difference between measurements made in the specimens and images on the `measurement dataset`. Vertical lines indicate the line *x =* 0, in which the difference between the measurements is zero. Each panel also shows the p-values for testing the hypothesis that the median difference between image and specimen measurements is zero.

**Figure 5:**
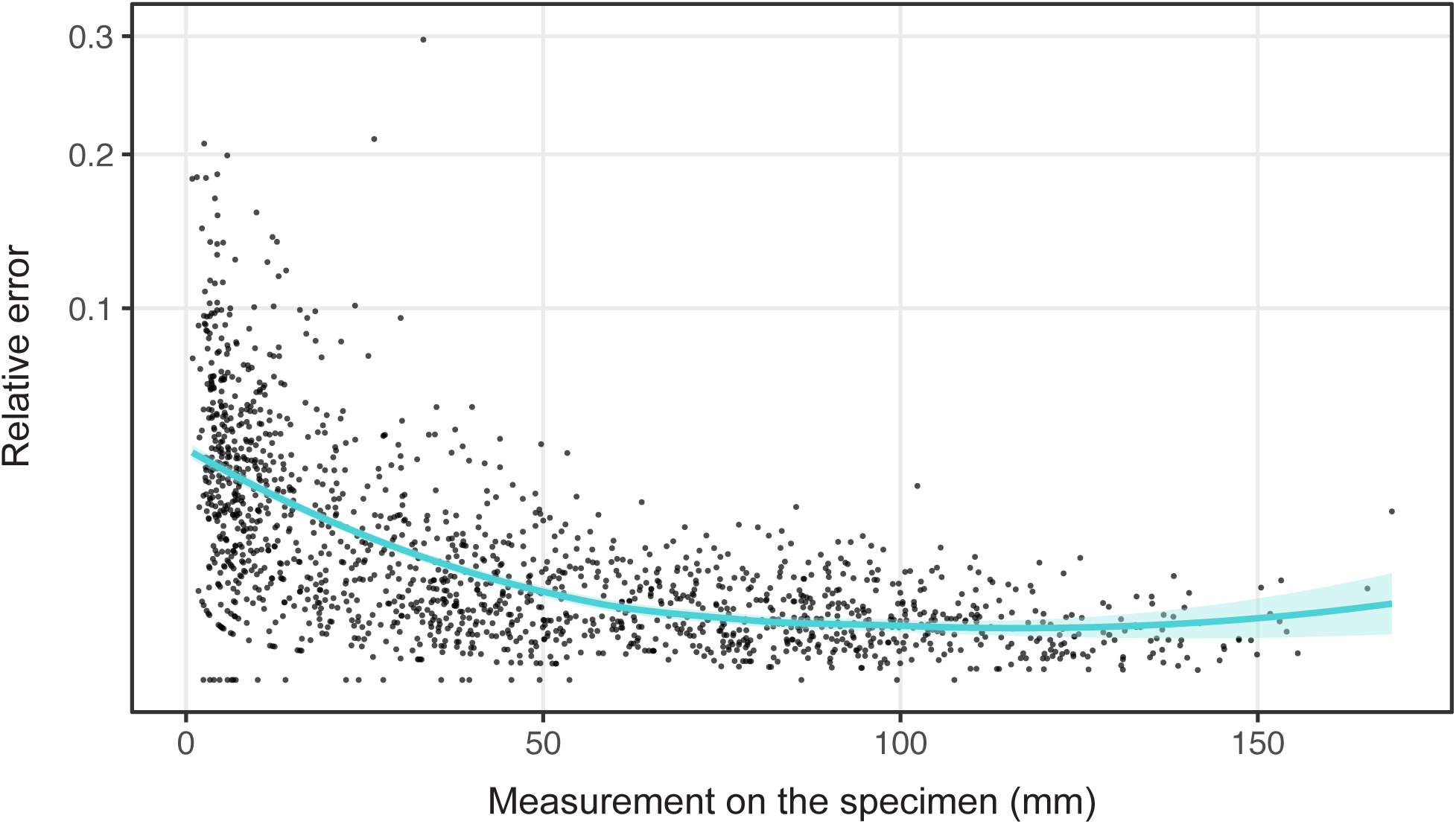
Relationship between the relative error of the measurements (i.e., the absolute difference between image and specimen measurements divided by specimen measurement) and the magnitude of each variable. Each dot represents a different observation from a variable in the `measurement dataset`. The curve indicates the mean trend, along with its standard error. *Y* -axis is displayed in a square root scale.

All but two `synthetic datasets` (E2015, T2008) include missing data. The total proportion of missing observations, percentage of variables with at least 50% of observations, and variables that were impossible to observe from images (Table 1) vary between `synthetic datasets`, but at least one of these classes lacks more than 25% of observations for most datasets. `Synthetic datasets` also differ in proportions of missing data by type of variable (continuous, categorical, discrete, or ratio; Table 2), with continuous variables being more commonly absent. Even though patterns of missing data may differ between datasets, correlation coefficients indicate that drops in accuracy are more related to absences spread in the dataset than to complete lack of a particular variable (Table 3).

**Table 1:** Proportion of missing data in each dataset according to three criteria: total percentage of missing data (total); percentage of variables with at least 50% of missing observations (at least 50%); percentage of variables impossible to score from images (100%).

| Dataset | total % | at least 50% | 100% |
| --- | --- | --- | --- |
| A2013 | 50% | 56.25% | 0% |
| A2008 | 44.39% | 45.12% | 19.51% |
| A2009 | 69.17% | 100% | 8.33% |
| B2018 | 37.5% | 35.71% | 35.71% |
| B2015 | 19.44% | 8.33% | 0% |
| E2015 | 0% | 0% | 0% |
| K2006i | 30.72% | 41.18% | 5.88% |
| K2006p | 30.72% | 41.18% | 5.88% |
| M1992 | 10% | 10% | 10% |
| P2005 | 42.97% | 43.75% | 12.5% |
| R2014 | 60.61% | 63.64% | 45.45% |
| S2012 | 43.56% | 44% | 24% |
| T2008 | 0% | 0% | 0% |

**Table 2:** Percentage of missing data by type of variable for each data set used to build the masks with missing data pattern.

| Dataset | Continuous (mm) | Categorical | Discrete | Ratio |
| --- | --- | --- | --- | --- |
| A2013 | 50% | - | - | - |
| A2008 | 39.39% | 53.33% | 52.73% | 37.65% |
| A2009 | 70% | - | 60% | - |
| B2018 | 50% | 16.67% | 0% | - |
| B2015 | 19.44% | - | - | - |
| E2015 | 0% | - | - | - |
| K2006i | 46.67% | - | 13.89% | 0% |
| K2006p | 46.67% | - | 13.89% | 0% |
| M1992 | 20% | 0% | 0% | - |
| P2005 | 32.81% | 56.25% | 43.75% | - |
| R2014 | 51.04% | - | 58.33% | 100% |
| S2012 | 41.11% | 56.94% | 31.75% | - |
| T2008 | 0% | - | 0% | - |

**Table 3:** Spearman correlation coefficients and their respective p-values for comparing the relationship between the accuracy in the estimation of each principal component/the PCA similarity (according to the “M” scheme) and the percentage of missing observations. total% - total percentage of missing data in the dataset; at least 50% - variables with at least 50% of missing observations; 100% - variables impossible to score from images.

| PCA Component | total % | at least 50% | 100% |
| --- | --- | --- | --- |
| 1 | -0.45 (0.12) | -0.48 (0.10) | -0.15 (0.63) |
| 2 | -0.78 (<0.01) | -0.79 (<0.01) | -0.51 (0.08) |
| 3 | -0.65 (0.02) | -0.66 (0.01) | -0.41 (0.16) |
| 4 | -0.67 (0.01) | -0.68 (0.01) | -0.30 (0.32) |
| PCA Sim | -0.73 (<0.01) | -0.73 (<0.01) | -0.78 (<0.01) |

The analyses of `original` and `synthetic datasets` (Fig. 6) show that the results of principal component analyses based on images are affected by noise and missing data. Nonetheless, noise effect is almost non-existent, as shown by values near 100% for both PCA similarity and PC correlation for the first and second principal components. On the other hand, accuracy drops in face of missing data, and particularly due to the joint effect of noise and missing data.

**Figure 6:**
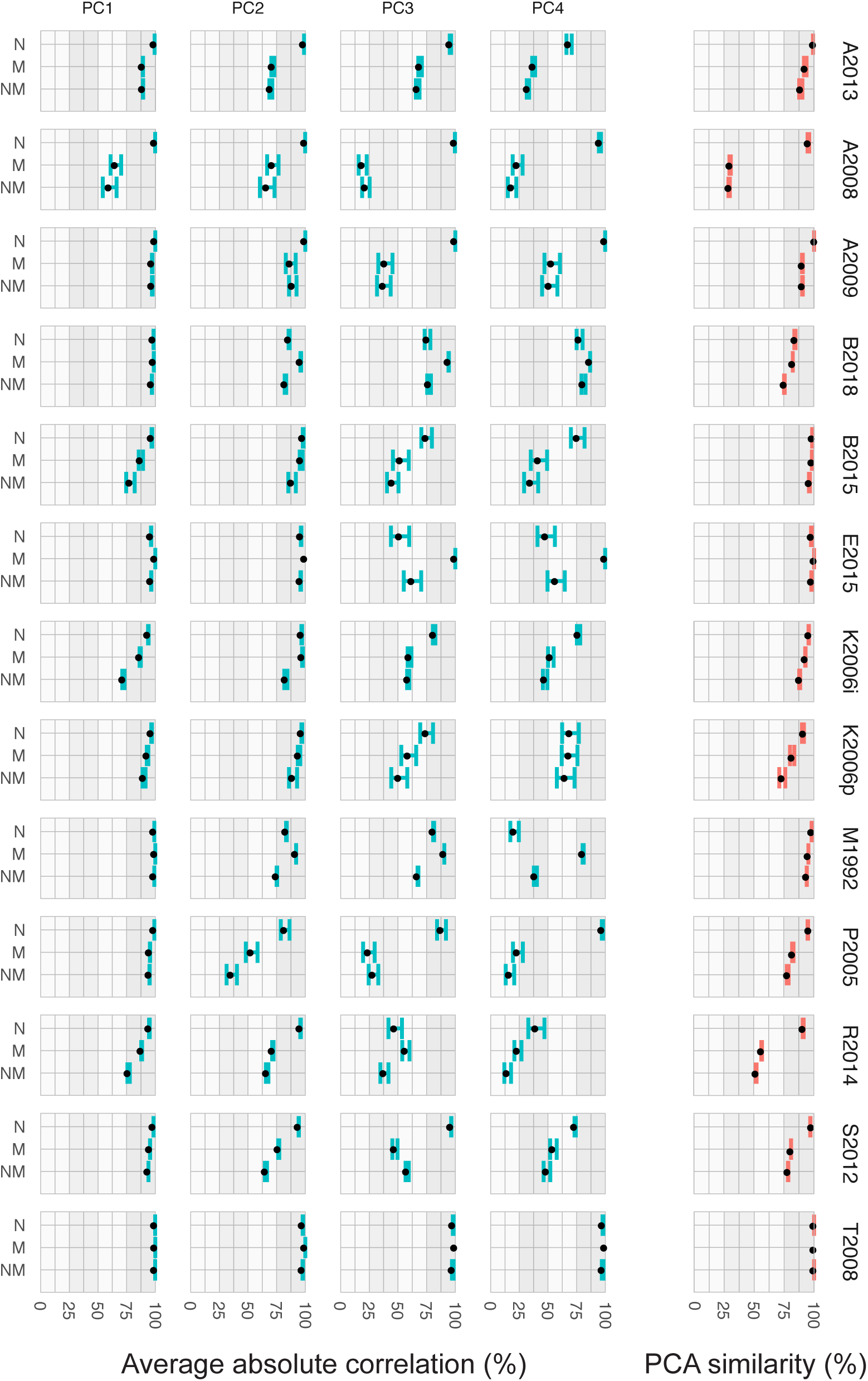
(Left) Accuracy (and 95% confidence intervals) in estimating the first four principal components using images (`synthetic datasets`). (Right) PCA Similarity between `synthetic datasets` versus `original`. 100% values equal maximum accuracy for both analyses. N - `synthetic dataset` with noise on the same level as those observed in Figure 3. M - `synthetic dataset` without variables that can not be acquired from images. NM - `synthetic dataset` with both noise and missing data.

## 4 Discussion

Here we used a set of leaf measurements and legacy morphometric studies to evaluate the precision and accuracy of morphological data gathered from images of herbarium specimens. We found image measurements to be highly precise and accurate, even though there is a small drop in accuracy for smaller traits. On the other hand, missing observations may decrease the accuracy of image-based morphological datasets. Below we discuss the problems, advantages, and consequences of collecting morphological data from digitized specimens.

### 4.1 Measurement precision and accuracy

Measurements taken from digitized herbarium specimens are as precise as the ones made on actual specimens. In fact, image measurements are in the confidence limits of being more precise (Fig. 2). An increase in precision has been seen for other imaging systems, as micro-computed tomography (Simon and Marroig, 2015). Thus, as we have focused only on leaves of particular species, it is possible analyses including a wider diversity of morphological variables to find the same for images of herbarium specimens.

Image measurements are also accurate, as indicated by their high similarity to specimen measurements (Fig. 3) and overall lack of bias (Fig. 4). Agreement between data gathered from images and specimens was also seen for the extraction of morphological features using artificial intelligence, which compared measurements taken by computers and researchers (Corney, Clark, Tang and Wilkin, 2012). Similar results for different organisms and imaging methods (Simon and Marroig, 2015; Aldridge et al., 2005) reinforce that high-confidence measurements can be obtained from digitized specimens. Nonetheless, we have also evidenced particular issues in the accuracy of image-based measurements and overall data acquisition.

### 4.2 Noise

The first problem with image use is a drop in measurement accuracy as variables scale down (Fig. 4). Measurements of smaller features, particularly those below 30–40 mm (Fig. 5), differ more between images and specimens than do larger structures. This association between accuracy error and scale was to be expected, as measurement precision lowers for smaller traits (Fig. 2). Such a bias in precision is not limited to images and is seen in our specimen measurements (Fig. 2), as well as for bird bones (Yezerinac et al., 1992) and *Drosera* (Droseraceae) leaves (Hoyo and Tsuyuzaki, 2013), for example. In this context, our results do not mean images are worse measurement sources. However, while we can increase precision and accuracy by studying specimens with different magnification tools, zooming images is restricted by resolution. Image resolution may not be a problem for high quality images (e.g. JStor Global Plants; https://plants.jstor.org/), but as digitization methods vary (Tulig et al., 2012; Takano et al., 2019; Sweeney et al., 2018), databases will differ in how they are prone to measurement errors.

Even though measurement errors will always be present, their effect is not strong. Our results show that datasets compiled from images and specimens are essentially the same (Fig. 6), especially as indicated by high PCA similarity (Yang and Shahabi, 2004; Melo et al., 2015). It is possible our results are biased, as most variables in our legacy datasets fall outside the range of sizes we found to be problematic for plant specimens (below 30–40 mm). Thus, researchers working on smaller organs or with small organisms should bear this potential bias in mind. Nonetheless, as seen for other taxa and methods using image-based measurements (e.g. Bruner, Emiliano and Costantini, David and Fanfani, Alberto and Dell’Omo, Giacomo, 2005; Corney, Tang, Clark, Hu and Jin, 2012; Corney, Clark, Tang and Wilkin, 2012; Chang and Alfaro, 2016), specimens two-dimensional images are good sources of metric data. Of course, this will only hold for traits that can be observed in the images.

### 4.3 Missing data

Another problem in our synthetic databases was the pervasive presence of missing data (Table 1), particularly of continuous variables (Table 2). As discussed above, these absences are likely due to image resolution issues, which hamper the observation of small features. Nonetheless, the extent of this problem is particular to each dataset. For example, the only two datasets with no missing data (T2008 and E2015; 1) also include variables that are smaller than 30 mm. At the same time, larger organs that were concealed during specimen preparation or due to their overlapping nature (e.g. petals usually hide both androecium and gynoecium) can not be observed and, thus, bias data collection. Feature-related bias is also seen in automatic trait recognition in which organ identity affects accuracy (Younis et al., 2018). Similar examples come from our datasets. If Trovó et al., 2008 had investigated the morphology of flowers (usually smaller than 2mm in Eriocaulaceae), it would not be possible to reproduce their data solely with images. This is the case with the T2013 dataset (Alcantara et al., 2013), in which the main source of missing data are organs concealed within the Bignoniaceae tubular flowers. Even though we can manipulate specimens to expose such hidden features, the same can not be done with herbarium images.

Indeed, missing data lower PC correlation and PCA similarity (Fig. 6). This trend is reinforced by the negative correlation between the proportion of missing data in `synthetic dataset` and PCA similarity or PC estimation (Table 3). Accuracy drops are stronger for the second, and especially for the third and fourth PCs. Although poor estimation of third and fourth PCs may not be a big problem, inaccuracy in estimation of the first two components and overall multivariate space, expressed in PCA similarity, points to significant differences between `original` and `synthetic datasets`. These differences could bias biological interpretations (Yezerinac et al., 1992) of studies relying on acquisition of morphological data, such as phenotypic evolution, disparity, modularity, phylogenetic inference, and others (e.g. Wiens, 2003; Zaragüeta-Bagils and Bourdon, 2007; Catalano et al., 2010; Klingenberg and Gidaszewski, 2010; Vasconcelos et al., 2018; Castiglione et al., 2019; Dellinger et al., 2019; Gallaher et al., 2019; Guillerme and Cooper, 2018).

### 4.4 Joint effect of measurement noise and missing data

Above, we analyzed the individual effects of measurement noise and missing data over dataset accuracy. However, most real datasets likely will include both issues. Our results indicate they interact, further dropping PC correlation and PCA similarity (Fig. 6). For a few cases in which noise is stronger than missing data (e.g. third and/or fourth PCs in B2018, and M1991), the joint effect improves component estimation, likely by removing noisy variables. Apart from those exceptions, the drop in accuracy when both noise and missing data are present may affect biological interpretations (see above). Particularly, rates of false positive and false negative conclusions may increase for study systems with low internal variation, which is common in comparisons of closely related taxa (Yezerinac et al., 1992).

Although the joint effect of noise and missing data may be serious, its influence will vary as datasets differ in the amount of missing data and presence of noise, which are linked to traits of interest size, image resolution, and organ overlap. Hence, before relying solely on morphological data compiled from digitized specimens, researchers should first test how image use affects their particular studies. For some cases, the overall accuracy we found in our analyses would be problematic (Fig. 6). For others, this accuracy would be just fine.

### 4.5 Consequences for image-based morphological data

When compiling morphological data from images of herbarium specimens, measurement noise is not a problem, particularly if variables are outside the size range affected by resolution. Hence, even though resolution may vary among digitization methods (Tulig et al., 2012; Takano et al., 2019; Sweeney et al., 2018), measuring them instead of specimens is an effective, less expensive (no need for traveling or loans), and, more important, reliable method for data acquisition.

On the other hand, missing data has a higher impact on data collection. This impact, however, will vary between different groups, particular morphologies, and specific research questions. For the cases in which missing data is relevant, researchers can improve data quality with particular strategies, used individually or in combination: 1) complete data with imputation methods; 2) exclude problematic variables or observations; 3) use specimens to fill gaps.

Data imputation methods estimate missing values in a dataset based on a variable’s internal variance, or on its associations to other variables in the dataset (Frane, 1976; Rubin, 1976; Little and Rubin, 2019). Here, for example, our PCA analyses used a relatively simple imputation method (Roweis, 1998). Although not free of issues (Little and Rubin, 2019), data estimation methods are powerful tools to improve results without compromising biological meaning (König et al., 2019).

Data imputation, however, will be inaccurate in cases where almost all observations of a given variable are missing. In this situation, it is sometimes better to exclude the variables from the analysis, particularly if it is highly correlated to other variables (Hemel et al., 1987). This way, variable exclusion is less likely to impact analyses or at least will have lower impact, as seen for the smaller negative correlation of fully absent variables to PC estimation and PCA similarity (Table 3). On the other hand, depending on the reason why a feature is missing, variable exclusion may bias results (Little and Rubin, 2019). Thus, it is important to understand the performance of data imputation and data exclusion when dealing with missing observations (Musil et al., 2002; Saunders et al., 2006; Horton and Kleinman, 2007; Enders, 2008).

If variable exclusion and data imputation are not feasible for a given case, such as a complete lack of observations on an important variable, researchers can use specimens to fill gaps in the dataset. Although not as practical as the other options, this approach can at least speed up work inside museums. When access to collections is limited, it is possible to combine examination of a minimum number of specimens and data imputation. These options exemplify how there is no single solution to circumvent missing data issues.

A more powerful approach to limit missing data – and further lower measurement noise – is to expand digital collections to include images with different magnifications, images exposing structures commonly hidden, or images taken from multiple angles. Increasing magnification and exposing hidden traits are complex and time-consuming but could be achieved if researchers fed back to institutions the images obtained while studying the specimens. Multi-camera imaging is particularly helpful for zoological specimens (Tegelberg et al., 2014; Price et al., 2018; Hereld et al., 2017; Ströbel et al., 2018; Hereld and Ferrier, 2019), as it allows proper visualization of traits that are hidden or distorted on images in a single view. Combining multi-view imaging and production lines (Tegelberg et al., 2014) is one efficient option to generate imagery that supports both 3D reconstructions (Hereld et al., 2017; Ströbel et al., 2018; Hereld and Ferrier, 2019) and accurate morphological data acquisition from the original two-dimensional images themselves, as shown here.

Despite particularities discussed above, digitization opened new frontiers for image-based phenotyping. Up to now, the bulk of research using specimen images focused on species identification (e.g. MacLeod, 2007; Cope et al., 2012; Wang, Lin, Ji and Liang, 2012; Wang, Ji, Liang and Yuan, 2012; MacLeod and Steart, 2015; Unger et al., 2016; Remagnino et al., 2016; Carranza-Rojas et al., 2017), or on shape analysis of particular organs (e.g. Corney, Clark, Tang and Wilkin, 2012; Smith and Kriebel, 2018; Reginato and Michelangeli, 2016). For studies focused on individual organs, datasets usually included only images of the focal traits. Now, the millions of images of herbarium specimens available make a wider diversity of phenotypes freely accessible, an accomplishment soon to be followed by zoological collections. In turn, easy access to specimen images supports data collection with crowdsourcing, a relatively fast and cheap way to obtain high-quality data (Chang and Alfaro, 2016; Zhou et al., 2018; O’Leary et al., 2018). Moreover, crowdsourced datasets may also aid the improvement of automatic extraction methods (Burleigh et al., 2013; Zhou et al., 2018), or be used in parallel with them to improve data collection. For example, while machines still can not differentiate overlapped plant organs (Gehan and Kellogg, 2017), an automatic approach can be complemented by human input, as we can easily identify and overcome overlapping issues. Independently of particular data collection strategies and of the need to be careful with missing observations, we reinforced here that digitized specimens are good sources of morphological data.

### 4.6 A note on data sharing

One of the main advantages of collecting data from digitized specimens is their free availability in digital repositories. It is only fair that sharing extends to data as well. However, at least for plant morphometric studies, this is not frequently done. For example, our `original dataset` came from just a few authors, who shared raw observations at publication time or kindly replied to our data requests. A much larger number of studies, however, presented only tables with mean and standard deviation values, which do not allow data reuse or verification.

If we aim to build comprehensive datasets, we have to shift this trend and increase the use of tools such as MorphoBank (https://morphobank.org; O’Leary and Kaufman, 2011) to share morphological data. Data sharing improves scientific work, and reduces redundant data collection, particularly if data is strongly integrated with ontologies (Deans et al., 2015) and linked to vouchers and scientific names in curated databases (Bruneau et al., 2019).

## 5 Conclusion

Measuring digitized specimens is a reliable data collection method, as demonstrated by a high correlation between measurement from specimens and images, even though resolution introduces measurement errors on small scales. At the same time, as it is impossible to manipulate images, particular features can not be observed and will be missing from image-based datasets. Even though there are some options to reduce the effects of missing data on analyses, the impact of this more serious problem has to be evaluated for each case.

Overall, we showed that accurate morphological data can be sourced from two-dimensional specimen images. For now, the millions of digital specimens (plus data on their distribution, ecology, and phenology) already available for plants (Le Bras et al., 2017; Soltis, 2017; Willis et al., 2017) put Botany at the front line of collective morphological data acquisition. As digitization expands to other groups, we will be able to merge big-data on geographical distribution, genomes, and morphology to unveil new knowledge on patterns and processes of phenotypic diversity and evolution.

## Acknowledgments

We are grateful to Abubakar Bello, Axel Poulsen and Inger Nordal, David Spooner and Mercedes Ames, Jeffrey Rose, Karol Marhold, Marcelo Trovó, Marek Slovák, Mariana Bünger, M. Montserrat Martínez-Ortega, Suzana Alcantara, and Vanessa Staggemeier for sharing their datasets, either at publication time or directly to us. We also thank Sara Mortara for initial help on research design and data collection, Kaori Nagata and Jon Wilcox for linguistic editing, as well as Rafaela C. Forzza, Paula Leitman, Fabiana L. R. Filard, and the RB staff for support during work at the herbarium. Anna Penna, Diogo Melo, Matheus F. Santos, and Monique N. Simon’s bright comments and suggestions helped us improve early versions of this work. Finally, we thank the editors and reviewers for their thoughtful suggestions. RI was supported in part by FAPESP (2019/11321-9) and CNPq (306943/2017-4). The authors declare no conflict of interest.

## Data accessibility statement

All data and code used here are available from GitHub (DOI: 10.5281/zenodo.3924506). Leaf measurements are also individually available from MorphoBank (project 3764 http://morphobank.org/permalink/?P3764).

